# DOT1L modulates the senescence-associated secretory phenotype through epigenetic regulation of IL1A

**DOI:** 10.1101/2020.08.21.258020

**Authors:** Kelly E. Leon, Raquel Buj, Elizabeth Lesko, Erika S. Dahl, Chi-Wei Chen, Yuka Imamura, Andrew V. Kossenkov, Ryan P. Hobbs, Katherine M. Aird

## Abstract

Cellular senescence is characterized as a stable cell cycle arrest that can occur as a stress response associated with oncogenic activation, termed oncogene-induced senescence (OIS). Cells undergoing OIS acquire a unique microenvironment termed the senescence-associated secretory phenotype (SASP), which can be both beneficial and detrimental in a context-dependent manner. Additionally, senescent cells are characterized by robust changes in their epigenome. Here, we globally assessed the histone landscape of cells induced to senesce by oncogenic RAS and discovered a novel epigenetic regulatory mechanism of the key SASP regulator IL1A. OIS cells displayed increased di- and tri-methylation of histone H3 lysine 79 (H3K79me2/3), two active histone marks. Depletion of the H3K79 methyltransferase disruptor of telomeric silencing 1-like (DOT1L) during OIS resulted in decreased H3K79me2/3 occupancy at the *IL1A* gene locus, which corresponded to decreased IL1A mRNA and cell surface expression. Decreased expression and secretion of downstream cytokines without a change in senescence markers were also observed upon DOT1L depletion. Overexpression of DOT1L increased H3K79me2/3 occupancy at the *IL1A* locus and upregulated the SASP, indicating that DOT1L is both necessary and sufficient for SASP gene expression. Mechanistically, we found that STING, an essential mediator of SASP transcription, is upstream of DOT1L in the epigenetic regulation of the SASP. Together, our studies establish DOT1L as an epigenetic regulator of the SASP whose expression is uncoupled from the senescence-associated cell cycle arrest, providing a potential strategy to inhibit the negative side effects of senescence while maintaining the beneficial inhibition of proliferation.

## Introduction

Cellular senescence is defined as a stable cell cycle arrest that can occur due to multiple stimuli, such as oncogenic stress (Serrano et al., 1997). Although the induction of senescence due to oncogene activation (termed oncogene-induced senescence: OIS) can result in tumor suppression, it may also result in tumor promotion and progression (Coppé et al., 2006; Ritschka et al., 2017; Sparmann and Bar-Sagi, 2004). One of the hallmarks of senescence is the senescence-associated secretory phenotype (SASP), a unique microenvironment composed of pro-inflammatory cytokines, chemokines, and matrix metalloproteinases (MMPs) (Acosta et al., 2008; Coppe et al., 2008; Kuilman et al., 2008). While the SASP may enhance immune cell recruitment and clearance of senescent cells, it also has detrimental side effects, resulting in chronic inflammation that contributes to tumorigenesis and chemoresistance (Coppe et al., 2008). Therefore, further understanding how to restrict the negative paracrine effects of the SASP while maintaining the senescence-associated cell cycle arrested has implications in transformation and tumorigenesis.

Previous studies have demonstrated that the pro-inflammatory cytokines and chemokines of the SASP are transcriptionally regulated (Acosta et al., 2008; Kuilman et al., 2008). One key component of the SASP is interleukin 1 alpha (IL1A), which is thought to be one of the critical upstream regulators of SASP expression (Gardner et al., 2015; Ong et al., 2018; Orjalo et al., 2009; Wiggins et al., 2019). Indeed, cell surface IL1A expression is necessary for the positive feedback loop between nuclear factor kappa B (NF-κβ) activation and transcription of multiple cytokines such as interleukin 6 (*IL6*), interleukin 8 (*CXCL8*; encoding IL8), and interleukin 1 beta (*IL1B*) (Orjalo et al., 2009). While target of rapamycin (mTOR) has been implicated in translational regulation of *IL1A* (Laberge et al., 2015), less is clear about its transcriptional regulation especially since it seems to be in part upstream of NF-κβ (Orjalo et al., 2009). Furthermore, recent studies have demonstrated that the innate DNA sensing pathway cyclic GMP-AMP synthase (cGAS)- stimulator of interferon genes (STING) is an upstream regulator of the SASP (Gluck et al., 2017; Yang et al., 2017). Increased DNA damage caused by OIS and decreased nuclear lamin expression results in cytoplasmic chromatin fragments that activate cGAS-STING and the downstream effectors interferon regulator factor 3 (IRF3) and NF-κβ (Di Micco et al., 2006; Dunphy et al., 2018; Gluck et al., 2017; Mackenzie et al., 2017). Although cGAS-STING has been implicated in regulating the SASP during OIS, whether and how cGAS-STING affects the transcription of the key SASP regulator *IL1A* is unknown.

In addition to the SASP, another hallmark of senescence is a marked change in histone modifications (Chandra et al., 2012; Narita et al., 2003; Zhang et al., 2007). For instance, di- and tri-methylation of H3K9, repressive histone marks that are found in heterochromatin, are known to decrease proliferation-promoting genes during OIS (Narita et al., 2003). However, SASP genes are protected from heterochromatinization via HMGB2 (Aird et al., 2016), allowing their continued and increased transcription. Previous reports have demonstrated that active and repressive histone marks such as H3K4me3 and H3K27me3, respectively, affect multiple senescence phenotypes including the SASP (Capell et al., 2016; Ito et al., 2018; Shah et al., 2013). Another histone mark that has been implicated in senescence is H3K79 (Kim et al., 2012; Wang et al., 2010). Methylation of H3K79 is associated with active transcription (Wood et al., 2018). H3K79 is methylated by the lysine methyltransferase disruptor of telomeric silencing 1-like (DOT1L) (Feng et al., 2002), and recent reports indicate that the Jumonji C (JmjC) demethylases KDM2B and KDM4D are likely H3K79 demethylases (Jbara et al., 2017; Kang et al., 2018). Methylation of H3K79 through DOT1L has been implicated in contributing to the DNA damage response (Kari et al., 2019). Additionally, previous studies have linked decreased DOT1L and H3K79 methylation to cell cycle inhibition and senescence (Barry et al., 2009; Kim et al., 2012). However, whether DOT1L or the active histone mark H3K79 regulate the SASP is unknown.

Here, we found that the active marks H3K79me2/3 were enriched at the *IL1A* gene locus in OIS cells. Mechanistically, we determined that the H3K79 methyltransferase DOT1L is upregulated in OIS, and DOT1L is both necessary and sufficient for H3K79me2/3 occupancy at the *IL1A* locus, leading to subsequent activation of downstream SASP genes without altering other senescence phenotypes. Upregulation of DOT1L required STING, suggesting that DOT1L upregulation downstream of cGAS-STING is a feed forward loop to increase SASP gene expression. Together, we determined that DOT1L is a central mediator of IL1A and SASP, which is uncoupled from the senescence-associated cell cycle arrest.

## Results

### H3K79 di- and tri-methylation marks are increased at the *IL1A* locus

Senescent cells have robust changes in their epigenome, including marked alterations in histone modifications (Narita et al., 2003; Zhang et al., 2007). To globally determine changes in histone modifications during senescence, we performed unbiased epiproteomics in the classic model of OIS: HRAS^G12V^ overexpression (Serrano et al., 1997), hereafter referred to as RAS (**Fig. S1A-F**). Multiple histone marks were significantly altered, including H3K27me3 and H3K36me1, which have been previously published (Chandra et al., 2012; Ito et al., 2018) (**Fig. 1A and Table S1**). Interestingly, H3K79me2 and H3K79me3, active histone marks, were increased during OIS, while the unmodified form of H3K79 was significantly decreased (**Fig. 1B**). Increased H3K79me2 and H3K79me3 was also observed by western blotting (**Fig. 1C**). We therefore sought to understand the functional role of H3K79 methylation in the regulation of OIS. To determine the global occupancy of H3K79me3 during OIS, we performed chromatin immunoprecipitation followed by next generation sequencing (ChIP-Seq). As H3K79me3 is an active histone mark (Wood et al., 2018), we then cross-compared ChIP-Seq peaks with genes that increase with a fold change >1.5 and q-value (FDR) <0.25 in our RNA-Seq analysis. This resulted in 95 common genes (**Fig. 1D and Table S2**). Gene set enrichment analysis (GSEA) of these 95 genes indicated enrichment of a number of pathways related to the immune response (**Fig. 1D**). Therefore, we reasoned that H3K79me3 may regulate the senescence-associated secretory phenotype (SASP) (Chien et al., 2011b; Kuilman et al., 2008; Rodier et al., 2009). Upon further analysis of our ChIP-Seq data, we discovered that H3K79me3 was enriched at the *IL1A* locus (**Fig. 1E**). While we observed an increase in mRNA expression of classical SASP genes, *IL6, IL1B, and CXCL8* (**Fig. S1G**), that are known to be transcriptionally regulated (Aird et al., 2016; Buj et al., 2020; Capell et al., 2016; Orjalo et al., 2009), H3K79me3 was not enriched at these loci (**Fig. S1H**). As IL1A is a critical upstream regulator of the SASP (Gardner et al., 2015; Ong et al., 2018; Orjalo et al., 2009; Wiggins et al., 2019), these data suggest that H3K79me3 may be important for initiating the SASP via *IL1A* transcription. H3K79me2 and H3K79me3 have distinct histone patterns, with H3K79me2 preferentially at the promoter region and H3K79me3 within the gene body (Guenther et al., 2007) (**Fig. S1I**). ChIP-qPCR validated H3K79me2 binding within the promoter region of the *IL1A* locus (**Fig. 1F**) and H3K79me3 within the gene body (**Fig. 1G**). Increased H3K79me2/3 at the *IL1A* gene locus corresponded to an increase in its transcription and expression at the cell surface (**Fig. 1H-I**). Together, these data suggest that H3K79me2/3 play a role in SASP gene expression during senescence via transcriptional activation of *IL1A*.

**Figure 1.**
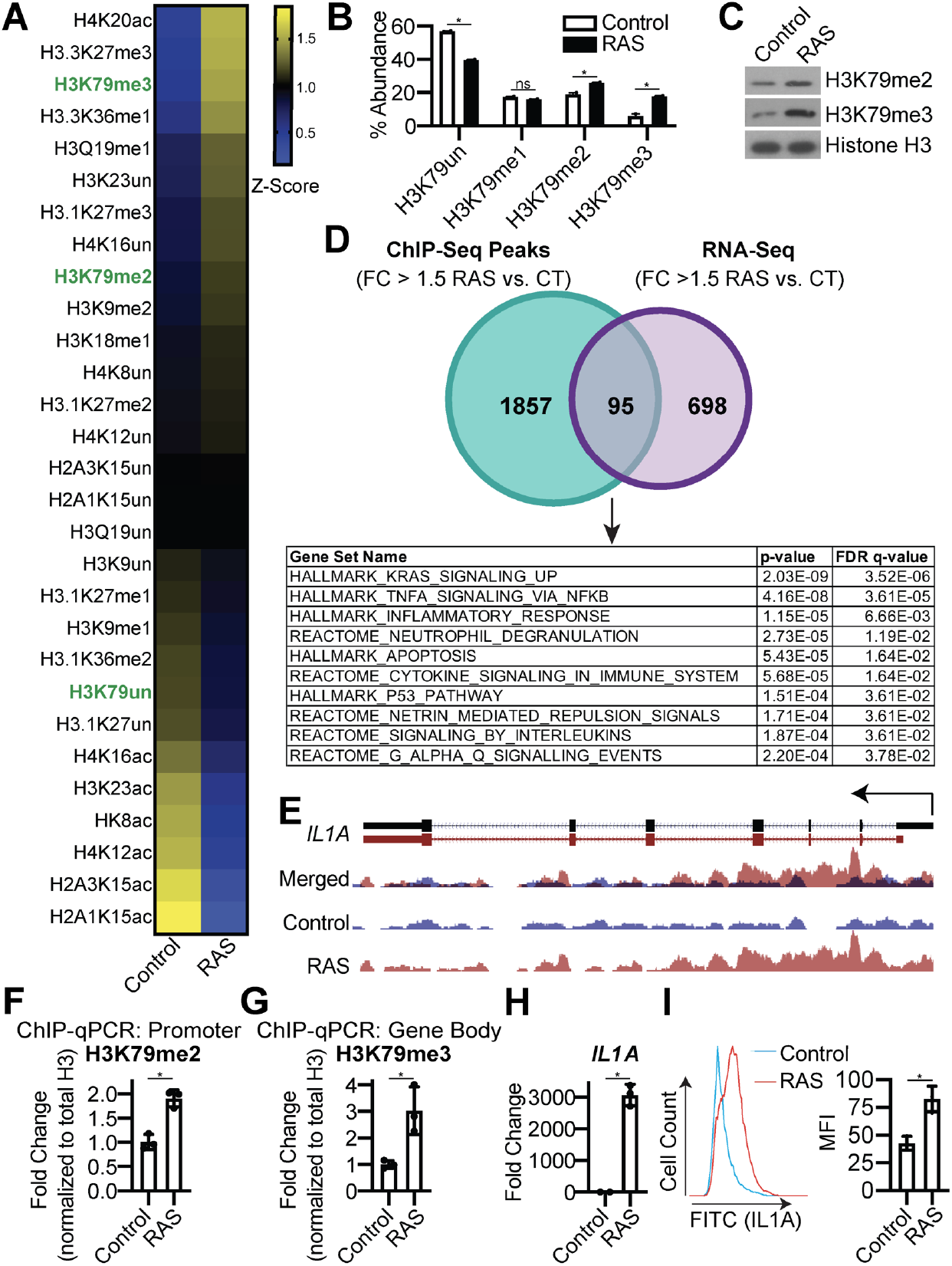
H3K79me2/3 are increased at the *IL1A* locus during oncogene-induced senescence. **(A-I)** IMR90 cells were infected with retrovirus expressing HRas^G12V^ (RAS) or empty vector control. See Figure S1 for timepoints. **(A)** Heatmap of unmodified, methylated, and acetylated histones determined using LC-MS/MS. H3K79 is marked in green. See Table S1 for raw data. **(B)** H3K79un (unmodified), H3K79me1, H3K79me2, and H3K79me3 percent abundance was determined by LC-MS/MS. Data represent mean ± SEM. *p<0.05. **(C)** H3K79me2 and H3K79me3 immunoblot analysis of chromatin fraction. Total histone H3 was used as a loading control. **(D)** Cross-referencing of H3K79me3 ChIP-Seq data and RNA-Seq data. A total of 1952 genes were bound by H3K79me3 (fold change >1.5 RAS vs. Control; Table S5) and a total of 793 genes were significantly increased by RNA-Seq [fold change >1.5 RAS vs. Control; q-value (FDR) <0.25; Table S6]. The 95 genes (listed in Table S2) that overlapped were subjected to gene set enrichment analysis. **(E)** H3K79me3 ChIP-Seq track at the *IL1A* gene locus. Blue indicates H3K79me3 binding in control cells, whereas red indicates H3K79me3 binding in RAS cells. **(F-G)** H3K79me2 binding to the *IL1A* promoter region **(F)** and H3K79me3 binding to the gene body **(G)** was determined by ChIP-qPCR and normalized to total histone H3 binding at the same site. One of three experiments is shown. Data represent mean ± SD. *p<0.05. **(H)** *IL1A* mRNA expression. One of five experiments is shown. Data represent mean ± SD. *p<0.001 **(I)** Cell surfacebound IL1A was determined by flow cytometry. MFI = median fluorescence intensity. One of four experiments is shown. Data represent mean ± SEM. *p<0.01.

### The H3K79 methyltransferase DOT1L is necessary for SASP gene expression

Next we aimed to determine the mechanism of increased H3K79 methylation in OIS. DOT1L is the only methyltransferase for H3K79 (Feng et al., 2002; Lacoste et al., 2002), while both KDM2B and KDM4D have been implicated as H3K79 demethylases (Jbara et al., 2017; Kang et al., 2018; Leon and Aird, 2019). RNA-Seq analysis of all three genes indicated that *DOT1L* is significantly upregulated in OIS cells, while there is no difference in either *KDM2B* or *KDM4D* expression (**Fig. S2A**). *DOT1L* was also upregulated in melanocytes induced to senesce by oncogenic BRAF^V600E^ and correlated with *IL1A* upregulation (**Fig. S2B-E**) (Pawlikowski et al., 2013), suggesting this is not a cell line or oncogene-specific phenotype. Consistent with the mRNA expression, we also observed an increase in DOT1L protein expression in OIS cells (**Fig. 2A**). To confirm DOT1L upregulation is necessary for H3K79 methylation in senescent cells, we knocked down DOT1L using two independent shRNAs in OIS cells (**Fig. 2B and Fig. S2F**). Knockdown of DOT1L decreased both H3K79me2 and H3K79me3 in OIS cells (**Fig. 2C**) and decreased occupancy of both DOT1L and H3K79me2 at the *IL1A* promoter (**Fig. 2D**) and both DOT1L and H3K79me3 occupancy within the gene body (**Fig. 2E**). This corresponded to decreased *IL1A* mRNA expression and IL1A expression at the cell membrane (**Fig. 2F-G**). We also observed a positive correlation between *DOT1L* and *IL1A* expression in mouse papillomas treated with DMBA/TPA, a known inducer of OIS and the SASP (Ritschka et al., 2017), suggesting this is phenomenon also occurs *in vivo* (**Fig. S2G**). Consistent with the idea that IL1A is the upstream regulator of the SASP, inhibition of H3K79 methylation at the *IL1A* locus by knockdown of DOT1L decreased transcription and secretion of other SASP factors (**Fig. 2H-I and Table S3**). The decrease in SASP was not due to abrogation of NF-κβ signaling or rescue of the senescence-associated cell cycle arrest (**Fig. S2H-O**). Consistently, expression of *CDKN2A* (encoding p16) and *CDKN1A* (encoding p21), two markers of senescence, did not decrease upon knockdown of DOT1L, suggesting H3K79me2/3 does not play a direct role in transcription of these genes during OIS (**Fig. S2P**). Additionally, decreased SASP expression was not due to a decrease in the DNA damage accumulation in DOT1L knockdown cells (**Fig. S2Q-R**). Pharmacological inhibition of DOT1L via treatment with the DOT1L inhibitor EPZ5676 also decreased H3K79me2 and H3K79me3 expression during OIS (**Fig. S2S-T**), which corresponded with decreased transcription of *IL1A* and other SASP factors (**Fig. S2U**). Consistent with genetic suppression DOT1L expression, pharmacological inhibition of DOT1L activity in OIS cells did not suppress senescence (**Fig. S2V-W**). These data demonstrate that DOT1L upregulation and H3K79me2/3 occupancy at *IL1A* is necessary for activating SASP gene expression during OIS.

**Figure 2.**
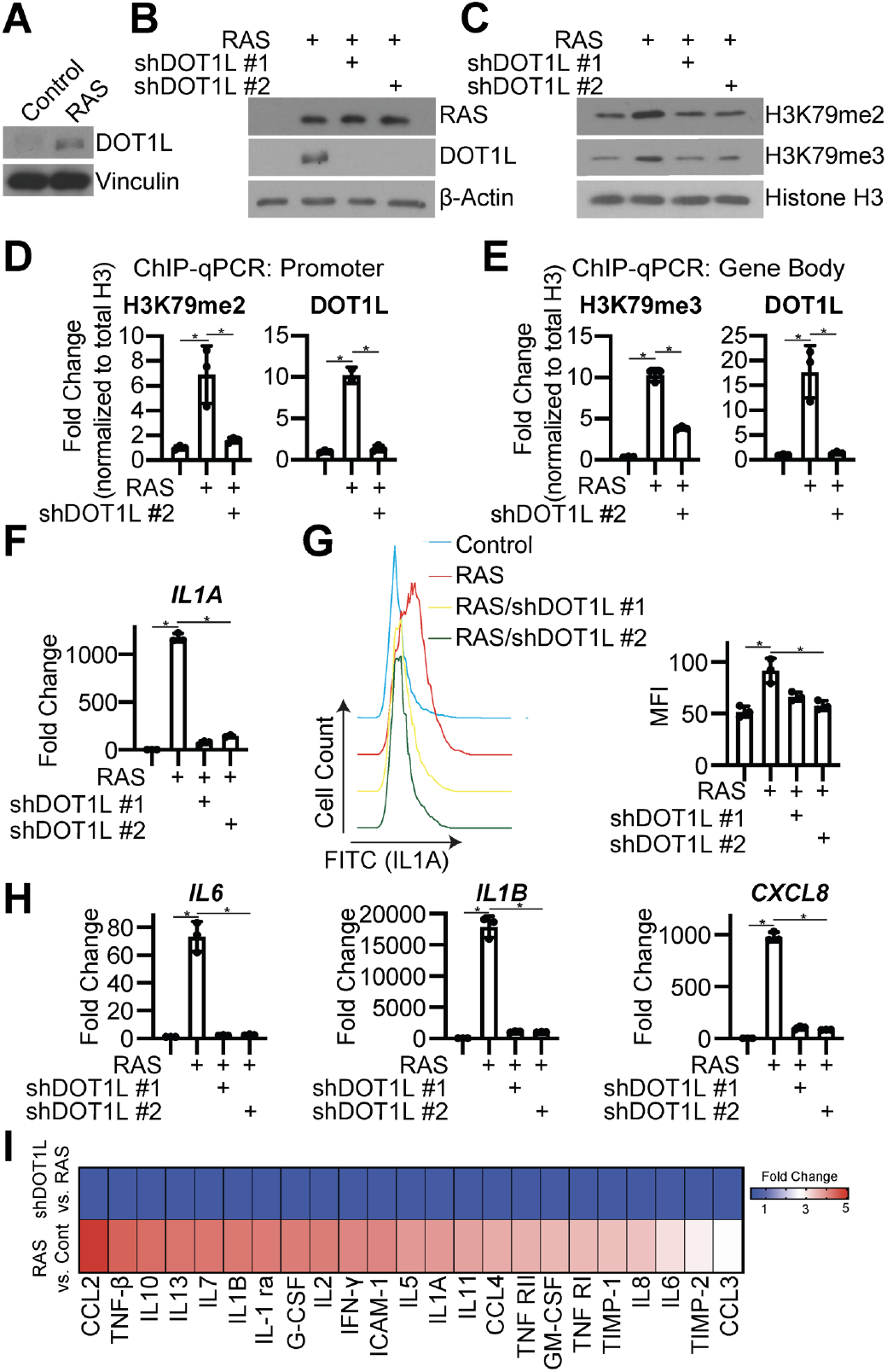
DOT1L is necessary for H3K79me2/3 at the *IL1A* locus and SASP expression. **(A)** IMR90 cells were infected with retrovirus expressing HRAS^G12V^ (RAS) or control. DOT1L protein expression was determined by immunoblotting. Vinculin was used as a loading control. One of five experiments is shown. **(B-I)** IMR90 cells were infected with retrovirus expressing HRAS^G12V^ (RAS) or control with or without shRNA to human DOT1L (shDOT1L). Details on timepoints are in Figure S1A. **(B)** RAS and DOT1L immunoblot analysis. β-actin was used as a loading control. One of three experiments is shown. **(C)** H3K79me2 and H3K79me3 immunoblot analysis on chromatin fractions. Histone H3 was used as a loading control. One of three experiments is shown. **(D-E)** H3K79me2 and DOT1L binding to the *IL1A* promoter region **(D)** and H3K79me3 and DOT1L binding to the *IL1A* gene body **(E)** was determined by ChIP-qPCR and normalized to total histone H3 binding at the same site. One of three experiments is shown. Data represent mean ± SD. *p<0.01. **(F)** *IL1A* mRNA expression was determined by RT-qPCR. One of five experiments is shown. Data represent mean ± SD. *p<0.01. **(G)** Cell surface-bound IL1A was determined by flow cytometry. MFI=median fluorescence intensity. One of four experiments is shown. Data represent mean ± SD. *p<0.01. **(H)** *IL6*, *IL1B*, and *CXCL8* mRNA expression was determined by RT-qPCR. One of five experiments is shown. Data represent mean ± SD. *p<0.01. **(I)** Secretion of SASP related factors was detected using an antibody array. Heat map indicates fold change. Raw data can be found in Table S3.

### H3K79 methylation drives SASP gene expression

Next we sought to determine whether DOT1L and H3K79me2/3 are sufficient for SASP induction. Towards this goal, we overexpressed DOT1L in normal IMR90 fibroblasts (**Fig. 3A and S3A**). Overexpression of DOT1L increased total H3K79me2 and H3K79me3 expression (**Fig. 3B**) and occupancy of both DOT1L and these active marks at the *IL1A* locus (**Fig. 3C-D**). Increased H3K79me2/3 occupancy at the *IL1A* locus corresponded to an increase in *IL1A* transcription, cell surface expression, and downstream SASP gene expression and secretion (**Fig. 3E-H and Table S3**). Overexpression of DOT1L did not increase markers of senescence or transcription of *CDKN2A* (encoding p16) or *CDKN1A* (encoding p21) (**Fig. S3B-G**). Importantly, increased SASP expression was also not due to increased DNA damage or decreased LMNB1 upon DOT1L overexpression as we did not observe an increase in DNA damage foci or depleted *LMNB1* in these cells (**Fig. S3H-K**). Together, these data indicate that DOT1L overexpression is sufficient to induce SASP expression through increased H3K79me2/3 at the *IL1A* locus.

**Figure 3.**
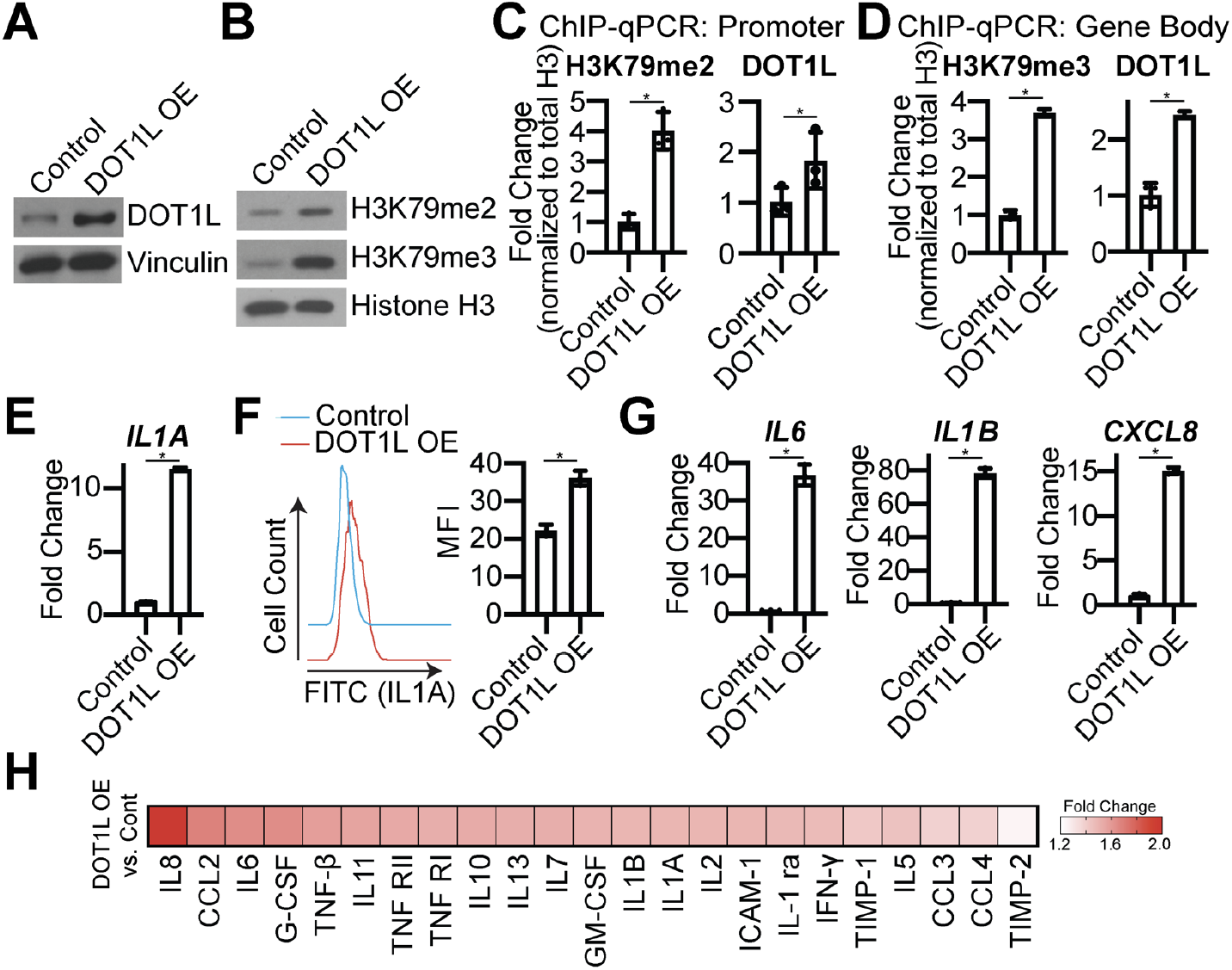
DOT1L overexpression increases H3K79me2/3 at the *IL1A* locus and is sufficient for SASP gene expression. **(A-H)** IMR90 cells were infected with retrovirus expressing human DOT1L or control. Cells were selected for 4 days with 1 μg/mL blasticidin. **(A)** DOT1L immunoblot analysis. Vinculin was used as a loading control. One of five experiments is shown. **(B)** H3K79me2 and H3K79me3 immunoblot analysis on chromatin fractions. Total histone H3 was used as loading control. One of three experiments is shown. **(C-D)** H3K79me2 and DOT1L binding to the *IL1A* promoter region **(C)** H3K79me3 and DOT1L binding to the *IL1A* gene body **(D)** was determined by ChIP-qPCR and normalized to total histone H3 binding at the same site. One of three experiments is shown. Data represent mean ± SD. *p<0.02. **(E)** *IL1A* mRNA expression was determined by RT-qPCR. One of five experiments is shown. Data represent mean ± SD. *p<0.006. **(F)** Cell surface-bound IL1A was determined by flow cytometry. MFI=median fluorescence intensity. One of four experiments is shown. Data represent mean ± SD. *p<0.001. **(G)** *IL6*, *IL1B*, and *CXCL8* mRNA expression was determined by RT-qPCR. One of five experiments is shown. Data represent mean ± SD. *p<0.0001. **(H)** Secretion of SASP related factors were detected by antibody array. Heat map indicates fold change versus control. Raw data can be found in Table S3.

### DOT1L is increased downstream of STING

Reports have demonstrated a role for cyclic GMP-AMP synthase (cGAS) in regulating the paracrine effects of the SASP via the stimulator of interferon genes (STING) (Dou et al., 2017; Gluck et al., 2017; Yang et al., 2017). Furthermore, a recent study implicated DOT1L in regulating the innate immune response (Chen et al., 2020). Therefore, we aimed to determine whether STING is upstream of DOT1L and H3K79 methylation during OIS. Towards this goal, we knocked down STING in OIS cells (**Fig. 4A-B**). STING knockdown decreased *DOT1L* expression in OIS cells (**Fig. 4C**). Consistently, we observed a decrease in expression of both H3K79me2 and H3K79me3 (**Fig. 4D**). As expected, STING knockdown decreased *IL1A* mRNA expression and transcription of downstream cytokines *IL6, IL1B*, and *CXCL8* (**Fig. 4E**). This was not due to rescue of senescence or changes in DNA damage as cells with STING knockdown maintained senescence markers and DNA damage accumulation (**Fig. S4A-G**). Rescue experiments using overexpression of DOT1L in STING knockdown OIS cells (**Fig. 4C**) abrogated the decrease in H3K79me2/3 and *IL1A* expression (**Fig. 4D-E**). Overexpression of DOT1L in STING knockdown OIS cells also rescued the expression of downstream cytokines *IL6, IL1B*, and *CXCL8* (**Fig. 4E**). These data implicate the cGAS-STING pathway in regulating the senescence microenvironment in part through DOT1L-mediated epigenetic control of *IL1A* and suggest that DOT1L expression participates in a feed forward mechanism to amplify SASP gene transcription (**Fig. 4F**).

**Figure 4.**
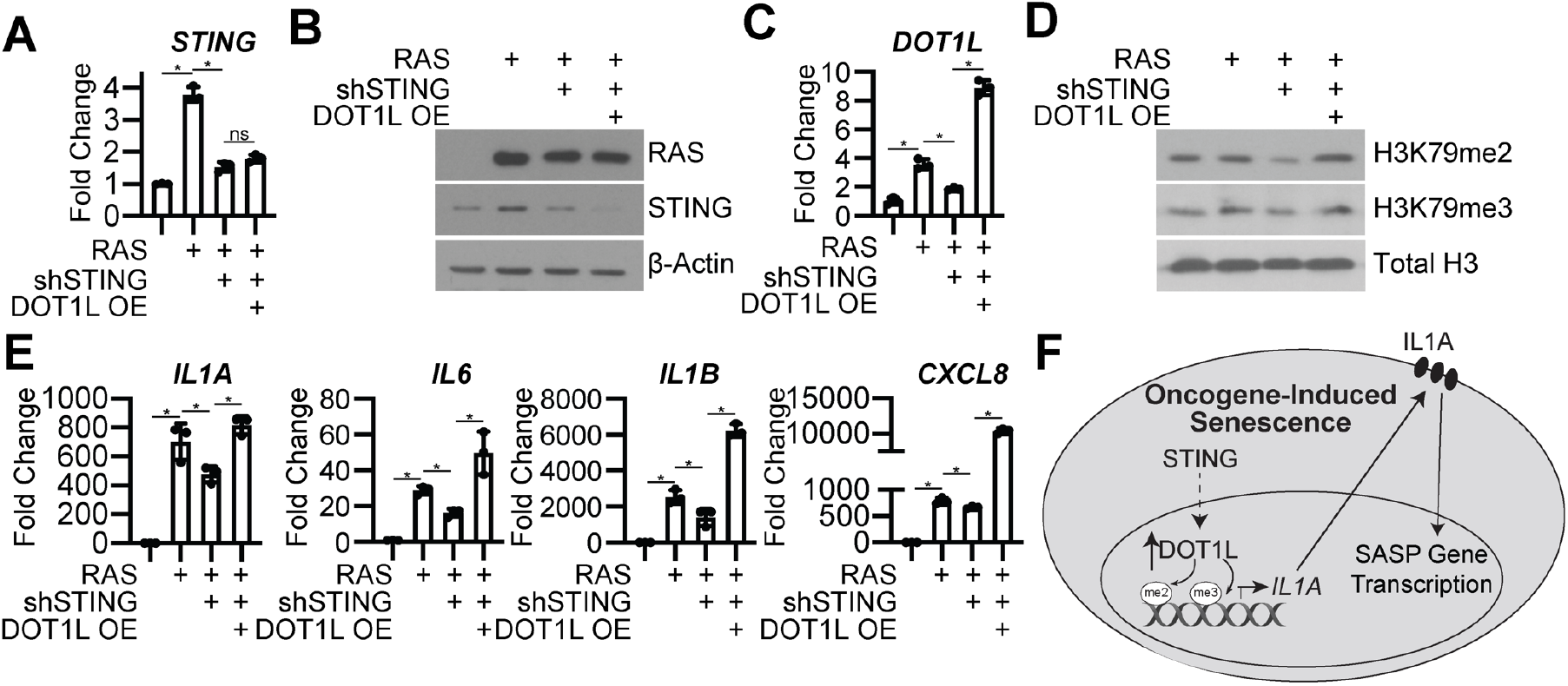
STING is necessary for DOT1L expression to promote the SASP. **(A-I)** IMR90 cells were infected with retrovirus expressing HRas^G12V^ (RAS) or control with or without lentivirus expressing an shRNA to human STING (shSTING) with or without overexpression of DOT1L (DOT1L OE). **(A)** *STING* mRNA expression was determined by RT-qPCR. One of three experiments is shown. Data represent mean ± SD. *p<0.0001. **(B)** RAS and STING immunoblot analysis. β-Actin was used as a loading control. One of three experiments is shown. **(C)** *DOT1L* mRNA expression was determined by RT-qPCR. One of three experiments is shown. Data represent mean ± SD. *p<0.0001. **(D)** H3K79me2 and H3K79me3 immunoblot analysis on the chromatin fraction. Total histone H3 was used as loading control. One of three experiments is shown. **(E)** *IL1A, IL6, IL1B*, and *CXCL8* mRNA expression was determined by RT-qPCR. One of five experiments is shown. Data represent mean ± SD. *p<0.001. **(F)** Proposed model of DOT1L-mediated SASP induction. STING induces DOT1L expression, which methylates H3K79 at the *IL1A* locus. IL1A is then transported to the cell surface, where it contributes to the feed forward mechanism of SASP induction through mediating SASP gene expression.

## Discussion

In the context of cancer, OIS is beneficial due to the repression of proliferation-promoting genes (Braig et al., 2005; Michaloglou et al., 2005; Serrano et al., 1997). However, although OIS results in a stable cell cycle arrest, cells actively transcribe and secrete a pro-inflammatory network of cytokines and chemokines, which may result in tumorigenesis (Cahu et al., 2012). Therefore, further mechanistic understanding of transcriptional regulation of the SASP is essential for developing strategies to prevent the detrimental effects of OIS while maintaining cells in a cell cycle arrested state. In this study, we found that OIS cells have increased expression of the H3K79 histone methyltransferase DOT1L, which is both necessary and sufficient for SASP expression through transcriptional regulation of the upstream SASP regulator *IL1A*. This phenotype required the STING pathway. Remarkably, while suppression of DOT1L decreased SASP gene expression and secretion, the cells maintained their senescent state. Together, these data indicate that while DOT1L expression contributes to the harmful expression and secretion of the SASP, it is uncoupled from the senescence-associated cell cycle arrest.

H3K79 is an active histone mark associated with transcriptional upregulation, DNA damage response, and cell cycle regulation (Feng et al., 2002; Giannattasio et al., 2005; Ng et al., 2003; Vakoc et al., 2006). DOT1L is the sole methyltransferase for H3K79 (Feng et al., 2002). We found that DOT1L and H3K79 methylation are increased in OIS cells (**Fig. 1 & 2**). Similarly, others have observed increased H3K79 methylation during OIS (Chicas et al., 2012) and increased H3K79 methylation at telomeres in yeast aging models (Rhie et al., 2013). Indeed, DOT1, the yeast homolog of DOT1L, suppresses senescence (Kozak et al., 2010), and DOT1L plays a protective role in UV-induced melanomagenesis (Zhu et al., 2018). In contrast, other studies have associated decreased H3K79 methylation and DOT1L expression with loss of proliferation and senescence (Barry et al., 2009; Jones et al., 2008; Kim et al., 2012; Nassa et al., 2019; Roidl et al., 2016; Song et al., 2020; Yang et al., 2019; Zhang et al., 2014). The discrepancy between these studies is not clear. It is possible that this is an inherent distinction between model systems. At least in the context of OIS versus replicative senescence in human cells, it is possible that oncogenic stress, which is generally more robust and associated with a hyperproliferative phase leading to replication stress and DNA damage (Aird et al., 2013; Di Micco et al., 2006), specifically leads to DOT1L upregulation and H3K79 methylation. Future studies will aim to reconcile these results.

We found that *DOT1L* is transcriptionally upregulated during OIS (**Fig. 2**). Previous studies have shown that the cytosolic DNA sensing pathway cGAS-STING is critical for SASP gene expression (Gluck et al., 2017; Yang et al., 2017), and knockdown of STING reduced *DOT1L* in OIS cells (**Fig. 4**), suggesting *DOT1L* is transcriptionally regulated downstream of cGAS-STING. A previous report linked the cGAS-STING pathway with toll like receptor 2 (TLR2) (Hari et al., 2019), and DOT1L is upregulated by activation of multiple TLRs (Chen et al., 2020). This suggests that in OIS, DOT1L may be upregulated downstream of the cGAS-STING-TLR2 axis. cGAS-STING is activated by cytosolic DNA during senescence due to increased DNA damage accumulation and decreased lamin B1 expression (Dou et al., 2017; Gluck et al., 2017). Interestingly, we found that overexpression of DOT1L alone increased H3K79me2/3 occupancy at *IL1A* and increased SASP gene expression without inducing DNA damage or altering LMNB1 expression (**Fig. S3**). DOT1L overexpression induced a modest fold change in SASP gene expression compared to OIS cells. This likely suggests that while DOT1L is sufficient to promote the SASP, its upregulation in OIS contributes to a feed forward mechanism downstream of DNA damage-mediated cGAS-STING-TLR2 activation (**Fig. 4F**). DOT1L is also a regulator of the DNA damage response, and H3K79 methylation promotes 53BP1 binding to site of double strand breaks (FitzGerald et al., 2011; Huyen et al., 2004). Interestingly, we did not observe marked changes in 53BP1 foci upon DOT1L knockdown (**Fig. S2**), again suggesting that while DNA damage is important for the feed forward mechanism, it is upstream of DOT1L.

H3K79 methylation is increased at the *IL1A* locus but not other SASP genes (**Fig. 1 and S1**). The question remains: how is DOT1L specifically recruited to the *IL1A* gene locus? DOT1L binding to chromatin is promoted by mono-ubiquitination of histone H2B on lysine 120 (H2BK120Ub) (Briggs et al., 2002; Kleer et al., 2002; McGinty et al., 2008), and it is possible that H2BK120Ub is elevated at the *IL1A* locus. Indeed, H3K4 methylation, which is also activated by H2BK120Ub (Kim et al., 2009) is upregulated by mixed lineage leukemia protein-1 (MLL1) at SASP gene loci (Capell et al., 2016). Additionally, DOT1L has multiple binding partners that could aid in its recruitment to particular loci including partner proteins (AF9, ENL, AF10 and AF17), MYC, and BAT3 (Cho et al., 2015; Wakeman et al., 2012; Wood et al., 2018), suggesting changes in expression or chromatin binding of these proteins may affect DOT1L recruitment to the *IL1A* gene locus. Finally, in yeast, DOT1 binding to chromatin is inhibited by Sir3 (Altaf et al., 2007; Fingerman et al., 2007), a human orthologue of sirtuins, whose expression and activity are often downregulated in senescence (Lee et al., 2019). Thus, it is possible that the loss of this inhibitory factor promotes DOT1L binding to the chromatin.

Numerous strides have been made towards clearing senescent cells in order to restrain the detrimental effects of cellular senescence. Indeed, selective targeting of senescent cells using genetic models or senolytics has clearly demonstrated that inhibition of the SASP in both cancer, aging, and other models is beneficial (Bussian et al., 2018; Chandra et al., 2020; Chang et al., 2016; Demaria et al., 2017; Jeon et al., 2017; Justice et al., 2019; Perrott et al., 2017; Suvakov et al., 2019; Wiley et al., 2018; Xu et al., 2015). While inhibitors of NF-κβ suppress the SASP, they also reverse the senescence-associated cell cycle arrest (Chien et al., 2011a), which would not be desirable in the context of cancer. Here we found that genetic knockdown or pharmacological inhibition of DOT1L using EPZ5676 suppressed the SASP while maintaining the senescence-associated cell cycle arrest. EPZ5676 is currently in clinical trials as a treatment for acute myeloid leukemia (Stein et al., 2018). Therefore, targeting the SASP through the use of DOT1L inhibitors may serve a novel therapeutic approach for age-related pathologies and cancer.

In conclusion, our study provides a new understanding of epigenetic regulation of the SASP via the H3K79 methyltransferase DOT1L. This layer of control likely reinforces the cGAS-STING-NF-κβ axis to transcriptionally increase expression of SASP genes in a feed forward pathway. As the SASP has been implicated in contributing to detrimental effects of senescence such as chemoresistance, tumor progression, and age-related pathologies, preventing SASP expression may be therapeutic in a wide range of diseases. Our findings provide rationale for targeting the H3K79 methyltransferase DOT1L to alleviate the harmful effects of the SASP while maintaining a senescent state.

## Materials and Methods

### Cells and Culture Conditions

Normal diploid IMR90 human fibroblasts were cultured in 2% O_2_ in DMEM (Corning; Cat.# 10017CV) with 10% FBS supplemented with L-glutamine, non-essential amino acids, sodium pyruvate, and sodium bicarbonate. IMR90s were used between population doubling #25-35. HEK293FT and Phoenix cells were cultured in DMEM (Corning; Cat.# 10013CV) with 5% FBS. Melanocytes were cultured in Melanocyte Growth Medium (Cell Applications, INC.; Cat# 135-500). All cell lines were cultured with MycoZap (Lonza; Cat.# VZA2032) and Penicillin-Streptomycin (Corning; Cat.# 30002Cl). All cell lines were routinely tested for mycoplasma as described in (Uphoff and Drexler, 2005).

### Plasmids and Antibodies

pBABE-puro-H-RAS^G12V^, MSCB-hDot1Lwt, and pHIV-BRAFV600E-mOrange2 were obtained from Addgene. pLKO.1-shDOT1L and pLKO.1-shSTING plasmids were obtained from Sigma-Aldrich with the following TRCN numbers: shDOT1L #1: TRCN0000020210; shDOT1L #2: TRCN0000020209; shSTING: TRCN00000160281. The following antibodies were obtained from the following suppliers: Anti Histone H3 tri methyl K79 (Abcam; Cat.# ab208189); Anti Histone H3 di methyl K79 (Abcam; Cat.# ab177184); Anti Histone H3 (Cell Signaling; Cat.# 14269); Anti DOT1L (Cell Signaling; Cat.# 77087); Anti STING (Cell Signaling; Cat.# 13647); Anti NF-κβ-p65 (Cell Signaling; Cat.# 8242S); Anti Phopho-NF-κβ-p65 (Cell Signaling; Cat.# 3033); Anti Cyclin A2 (Abcam; ab181591); Anti CDKN2A/p16INK4a (Abcam; Cat.# ab108349); Anti p53 (Calbiochem; Cat.# OP43); Anti Ras (BD Transduction Laboratories; Cat.# 610001); Anti BRAF (Santa Cruz; Cat.# sc-5284); Anti Vinculin (Sigma-Aldrich; Cat.# V9131); Anti Beta-Actin (Sigma-Aldrich; Cat.# A1978); Anti PML (Santa Cruz; Cat.# sc-966); Anti γH2AX (EMD Millipore; Cat.# 05-636); Anti 53BP1 (Bethyl Laboratories; Cat.# A300-272A); Anti IgG1 kappa (Thermo Fisher Scientific; Cat.# 50-186-16); Anti IL-1 alpha (Thermo Fisher Scientific; Cat.# 11-7118-81).

### Retroviral and Lentiviral Transduction and Infection

Phoenix cells (a gift from Dr. Gary Nolan, Stanford University) were used to package retroviral infection viruses. Retrovirus production and transduction were performed using the BBS/calcium chloride method as described in (Aird et al., 2013). HEK293FT cells were used to package lentiviral infection viruses using the ViraPower kit (Invitrogen, Carlsbad, California, USA) as per the manufacturer’s instructions.

IMR90 cells were infected with retrovirus and/or lentivirus as indicated in **Figure S1A**. IMR90 cells were infected with two rounds of pBABE-Control or pBABE-HRAS^G12V^. Cells were selected with 1μg/mL puromycin for 7 days. IMR90 cells were also infected with pLKO.1-shGFP (Control), pLKO.1-shDOT1L, or pLKO.1-shSTING where indicated and selected with 3μg/mL puromycin for 4 days. Alternatively, IMR90 cells were infected with two rounds of MSCB-hDot1lwt and selected with 1μg/mL blasticidin for 4 days. For the rescue experiment, IMR90 cells were initially infected with pBABE-Control or pBABE-HRAS^G12V^ followed by a simultaneous infection with MSCB-hDot1lwt and pLKO.1-shGFP or pLKO.1-shSTING. Cells were selected with puromycin and blasticidin for 7 days. Melanocyte cells were infected with pLKO.1-shGFP or pHIV-BRAFV600E-mOrange2 followed by collection for experiments 30 days post-infection.

### Mass Spectrometry Analysis of Histone Modifications

The cell pellet was resuspended in nuclear isolation buffer [15mM Tris-HCl (pH 7.5), 60mM KCl, 15mM NaCl, 5mM MgCl_2_, 1mM CaCl_2_, 250mM Sucrose, 1mM DTT, 1:100 Halt Protease Inhibitor Cocktail (Thermo Scientific), and 10mM sodium butyrate]. Nuclei were resuspended in 0.2M H_2_SO_4_ for 1 hour at room temperature and centrifuged at 4000 *x g* for 5 minutes. Histones were precipitated from the supernatant by the addition of TCA at a final concentration of 20% TCA (v/v). Precipitated histones were pelleted at 10,000 *x g* for 5 minutes, washed once with 0.1% HCl in acetone and twice with acetone followed by centrifugation at 14,000 *x g* for 5 minutes. Histones were air dried then resuspended in 10 μL of 0.1 M (NH)_4_HCO_3_ for derivatization and digestion according to (Garcia et al., 2007). Peptides were resuspended in 100 μL 0.1% TFA in water for LC-MS/MS analysis.

Multiple reaction monitoring was performed on a triple quadrupole (QqQ) mass spectrometer (ThermoFisher Scientific TSQ Quantiva) coupled with an UltiMate 3000 Dionex nano-LC system. Peptides were loaded with 0.1% TFA in water at 2.5 μl/minute for 10 minutes onto a trapping column (3 cm × 150 μm, Bischoff ProntoSIL C18-AQ, 3 μm, 200 Å resin) and then separated on a New Objective PicoChip analytical column (10 cm × 75 μm, ProntoSIL C18-AQ, 3 μm, 200 Å resin). Separation of peptides was achieved using solvent A (0.1% formic acid in water) and solvent B (0.1% formic acid in 95% acetonitrile) with the following gradient: 0 to 35% solvent B at a flow rate of 0.30 μl/minute over 45 minutes. The following QqQ settings were used across all analyses: collision gas pressure of 1.5 mTorr; Q1 peak width of 0.7 (FWHM); cycle time of 2 s; skimmer offset of 10 V; electrospray voltage of 2.5 kV. Monitored peptides were selected based on previous reports (Zheng et al., 2012; Zheng et al., 2013).

Raw MS files were imported and analyzed in Skyline software with Savitzky-Golay smoothing (MacLean et al., 2010). Automatic peak assignments from Skyline were manually confirmed. Peptide peak areas from Skyline were used to determine the relative abundance of each histone modification by calculating the peptide peak area for a peptide of interest and dividing by the sum of the peak areas for all peptides with that sequence. The relative abundances were determined based on the mean of three technical replicates with error bars representing the standard deviation.

### Western Blotting

Cell lysates were collected in 1X sample buffer (2% SDS, 10% glycerol, 0.01% bromophenol blue, 62.5mM Tris, pH 6.8, 0.1M DTT), boiled to 95°C for 10 minutes and sonicated. Chromatin samples were collected by trypsinizing and pelleting cells in 1X PBS at 3000rpm for 4 minutes. Cell pellets were washed with 1X PBS and pelleted again at 3000rpm for 4 minutes. Cell pellets were resuspended in a 10mM HEPES-KOH (pH 8.0), 10mM KCl, 1.5mM MgCL_2_, 0.34M sucrose, 10% glycerol, 1mM DTT, 0.1mM PMSF, and proteinase inhibitor cocktail buffer with a final pH of 7.5 (Buffer A). 0.1% Triton-X 100 was added to each sample and incubated on ice for 5 minutes. Samples were then centrifuged at 1300g for 4 minutes at 4°C. Following centrifugation, buffer was removed followed by addition of Buffer A and centrifugation at 1300g for 4 minutes at 4°C two additional times. Buffer A was completely removed and cell pellets were resuspended in 3mM EDTA (pH 8.0), 0.2mM EGTA, 1mM DTT, 0.1mM PMSF, and proteinase inhibitor cocktail buffer with a final pH of 8.0 (Buffer B) and incubated on ice for 30 minutes. Samples were then centrifuged at 1700g for 4 minutes at 4°C. Following centrifugation, buffer was removed followed by addition of Buffer B and centrifugation at 1700g for 4 minutes at 4°C two additional times. Buffer B was aspirated and half volume of 3X sample buffer and 1 volume 1X sample buffer was added to each chromatin pellet.

Total protein and chromatin concentrations were obtained using the Bradford assay. An equal amount of total protein or chromatin were resolved using SDS-PAGE gels and transferred onto nitrocellulose membranes. Membranes were blocked with 5% nonfat milk or 4% BSA in TBS containing 0.1% Tween-20 (TBS-T) for 1 hour at room temperature. Membranes were incubated overnight at 4°C in primary antibodies in 4% BSA/TBS and 0.025% sodium azide. Membranes were then washed 4 times in TBS-T for 5 minutes at room temperature followed by an incubation with HRP-conjugated secondary antibodies listed above. Membranes were washed 4 times in TBS-T for 5 minutes at room temperature and visualized on film with SuperSignal West Pico PLUS Chemiluminescent Substrate (Thermo Fisher, Waltham, MA).

### Chromatin Immunoprecipitation (ChIP) and ChIP-Seq

ChIP was performed as previously described in (Aird et al., 2016) using ChIP-grade antibodies described above. Immunoprecipitated DNA was analyzed by qPCR using iTaq Universal SYBR^®^ Green Supermix (BioRad; Cat.# 1725121). The following amplification conditions were used: 5 minutes at 95°C, 40 cycles of 95°C for 10 seconds and 30 seconds with 62°C annealing temperature. The assay ended with a melting-curve program: 15 seconds at 95°C, 1 minute at 60°C, then ramping to 95°C while continuously monitoring fluorescence. All samples were assessed in triplicate. Enrichment of H3K79me2, H3K79me3, and DOT1L was determined and normalized to total Histone H3. ChIP-qPCR primer sets are described in **Table S4**.

IMR90 cells induced to senescence by oncogenic RAS were prepared and analyzed through ChIP-Seq using anti-histone H3 tri methyl K79 (Abcam; Cat.# ab208189). ChIP-Seq libraries were created using Kapa HyperPrep Kit (Roche Sequencing and Life Science). The unique dual index sequences (NEXTFLEX^®^ Unique Dual Index Barcodes, BioO Scientific) were incorporated in the adaptors for multiplexed high-throughput sequencing. The final product was assessed for its size distribution and concentration using BioAnalyzer High Sensitivity DNA Kit (Agilent Technologies). The libraries were pooled and diluted to 3 nM using 10 mM Tris-HCl, pH 8.5 and then denatured using the Illumina protocol. The denatured libraries were loaded onto an S1 flow cell on an Illumina NovaSeq 6000 (Illumina) and run for 2X50 cycles according to the manufacturer’s instructions. De-multiplexed and adapter-trimmed sequencing reads were generated using Illumina bcl2fastq (released version 2.20.0.422). BBDuk (sourceforge.net/projects/bbmap/) was used to trim/filter low quality sequences using “qtrim=lr trimq=10 maq=10” option. Next, alignment of the filtered reads to the human reference genome (GRCh38) was done using bowtie2 (version 2.3.4.3) (Langmead and Salzberg, 2012). Only sequences aligning uniquely to the human genome were used to identify peaks. Peaks were called using MACS2 (version 2.1.1) and annotated using homer (version 4.10) by supplying GRCh38.78.gtf as the annotation file.

For the ChIP-Seq analysis, raw ChIP-Seq data reads were aligned against hg19 version of human genome using *bowtie* (Langmead et al., 2009) and *HOMER* (Heinz et al., 2010) was used to generate bigwig files and call significant peaks vs input using “-style histone” option. Peaks that passed 4 fold, FDR<5% threshold were considered. Normalized signals for significant peaks were derived from bigwig files using *bigWigAverageOverBed* tool from UCSC toolbox (Kent et al., 2010) with mean0 option. Fold differences between samples were then calculated with average input signal 0.4 used as a floor for the minimum allowed signal. Genes with transcripts body overlapping significant peaks were considered to be putatively regulated by the histone modification. Processed data can be found in **Table S5**.

### RNA-Seq

A total of 1 μg of RNA was used for cDNA library construction at Novogene using an NEBNext Ultra 2 RNA Library Prep Kit for Illumina (New England Biolabs; Cat.# E7775) as per the manufacturer’s instructions. Briefly, mRNA was enriched using oligo(dT) beads followed by two rounds of purification and fragmented randomly by adding fragmentation buffer. The first strand cDNA was synthesized using random hexamers primer, after which a custom second-strand synthesis buffer (Illumina), dNTPs, RNase H and DNA polymerase I were added to generate the second strand (ds cDNA). After a series of terminal repair, poly-adenylation, and sequencing adaptor ligation, the double-stranded cDNA library was completed following size selection and PCR enrichment. The resulting 250-350 bp insert libraries were quantified using a Qubit 2.0 fluorometer (Thermo Fisher Scientific) and quantitative PCR. Size distribution was analyzed using an Agilent 2100 Bioanalyzer (Agilent Technologies). Qualified libraries were sequenced on an Illumina NovaSeq 6000 Platform (Illumina) using a paired-end 150 run (2×150 bases). 20 M raw reads were generated from each library.

For RNA-Seq analysis, reference genome and gene model annotation files were downloaded from genome website browser (NCBI/UCSC/Ensembl) directly. Indexes of the reference genome was built using STAR and paired-end clean reads were aligned to the reference genome using STAR (v2.5). STAR used the method of Maximal Mappable Prefix(MMP) which can generate a precise mapping result for junction reads. STAR will count number reads per gene while mapping. The counts coincide with those produced by htseq-count with default parameters. And then FPKM of each gene was calculated based on the length of the gene and reads count mapped to this gene. FPKM, Reads Per Kilobase of exon model per Million mapped reads, considers the effect of sequencing depth and gene length for the reads count at the same time, and is currently the most commonly used method for estimating gene expression levels (Mortazavi et al., 2008). Differential expression analysis between two conditions/groups (two biological replicates per condition) was performed using the DESeq2 R package (1.14.1). DESeq2 provide statistical routines for determining differential expression in digital gene expression data using a model based on the negative binomial distribution. The resulting P-values were adjusted using the Benjamini and Hochberg’s approach for controlling the False Discovery Rate (FDR). Genes with an adjusted P-value <0.05 found by DESeq2 were assigned as differentially expressed. Processed data can be found in **Table S6**.

### Gene Set Enrichment Analysis

Common genes between ChIP-Seq and RNA-Seq (FC > 1.5 RAS vs. control) were analyzed using the Broad Institute Gene Set Enrichment Analysis (GSEA) webtool (https://www.gsea-msigdb.org/gsea/msigdb/annotate.jsp) for Hallmarks and Reactome. Following GSEA documentation, indications terms were considered significant when the FDR adjusted p-value (q-value) was <0.25.

### Quantitative RT-PCR (RT-qPCR)

Total RNA was extracted from cells using Trizol and DNase treated, cleaned, and concentrated using Zymo columns (Zymo Research, Cat.# R1013) as per manufacture instructions. Optimal density values of extracted RNA was obtained using NanoDrop One (Thermo Fisher Scientific). Only RNA with A260 and A280 rations above 1.9 was used. Relative expression of target genes listed in **Table S4** were analyzed using the QuantStudio 3 Real-Time PCR System (Thermo Fisher Scientific). All primers were designed using the Integrated DNA Technologies (IDT) tool (http://eu.idtdna.com/scitools/Applications/RealTimePCR/) (**Table S4**). A total of 25ng RNA was used in One-Step qPCR (Quanta BioSciences; Cat.# 95089) as per the manufacturer’s instructions. The following amplification conditions were used: 10 minutes at 48°C, 5 minutes at 95°C, 40 cycles of 10 seconds at 95°C and 7 seconds at the annealing temperature of 62°C. The assay ended with a melting-curve program: 15 seconds at 95°C, 1 minutes at 70°C, then ramping to 95°C while continuously monitoring fluorescence. All samples were assessed in triplicate. Relative quantification was determined normalizing to multiple reference genes (*MRPL9, PUM1, PSMC4*) using the delta-delta Ct method.

### Cell Surface Staining of IL1A

Cells were washed 2 times with 1X PBS and harvested using trypsin. Cells were pelleted by centrifugation at 1000rpm for 5 minutes at 4°C and then washed twice with 3% BSA/PBS. Cells were stained (10 μL/1.5×10^5^ cells) with anti-IL-1 alpha (Thermo Fisher Scientific; Cat.# 11-7118-81) and anti IgG1 kappa (Thermo Fisher Scientific; Cat.# 50-186-16) for 1 hour on ice. An unstained control was also incubated on ice for 1 hour. Cells were then pelleted by centrifugation at 1000rpm for 5 minutes at 4°C and washed with 3% BSA/PBS. Cell pellets were resuspended in 200 μL 3% BSA/PBS. Stained cells and unstained control cells were run on a 10-color FACSCanto flow cytometer (BD BioSciences). All data were analyzed using FlowJo Software.

### Cytokine Secretion Quantification

The human cytokine antibody array C1000 (RayBio; Cat.# AAH-CYT-1000-2) was used as per the manufacturer’s instructions to quantify secreted factors. Conditioned media was obtained from cells that were washed with 1X PBS and incubated in serum-free media for 48 hours. A 0.2 μm filter was used to filter conditioned media. Membranes were visualized on film. Individual spot signal densities were obtained using the ImageJ Software and normalized to cell number from which the conditioned media was obtained.

### Senescence-Associated Beta Galactosidase Activity Assay

Cells were seeded at an equal density 24 hours prior to assay. Cells were fixed for 5 minutes at room temperature using 2% formaldehyde/0.2% glutaraldehyde in 1X PBS. Cells were then washed twice with 1X PBS and stained with 40mM Na_2_H PO_4_, 150mM NaCl, 2mM MgCl_2_, 5mM K_3_Fe(CN)_6_, 5mM K_4_Fe(CN)_6_, and 1mg/mL X-gal staining solution overnight at 37°C in a non-CO_2_ incubator.

### Immunofluorescence

Cells were seeded at an equal density on coverslips and fixed with 4% paraformaldehyde. Cells were washed 4 times with 1X PBS and permeabilized with 0.2% Triton-X 100 in PBS for 5 minutes and then post-fixed with 1% paraformaldehyde and 0.01% Tween-20 for 30 minutes. Cells were blocked for 5 minutes with 3% BSA/PBS followed by incubation of corresponding primary antibody in 3% BSA/PBS for 1 hour at room temperature. Prior to incubation with secondary antibody in 3% BSA/PBS for 1 hour at room temperature, cells were washed 3 times with 1% Triton-X 100 in PBS. Cells were then incubated with 0.15μg/mL DAPI in 1X PBS for 1 minute, washed 3 times with 1X PBS, mounted and sealed. Images were all obtained using the Nikon Eclipse 90i microscope with a 20x/0.17 objective (Nikon DIC N2 Plan Apo).

### Colony Formation Assay

Cells were seeded at an equal density in 6-well plates and cultured for 14 days. Cells were then fixed using 1% paraformaldehyde/PBS and stained with 0.05% crystal violet. 10% acetic acid was used to destain and absorbance was measured at 590nm using a Spectra Max 190 spectrophotometer.

### DMBA/TPA Mouse Treatment

The shaved back skin of female wildtype FVB/NJ mice between 7 and 8 weeks of age was treated topically with a single dose of DMBA (7,12-Dimethylbenz(*a*)anthracene) in acetone (25 μg in 250 μL). Beginning one week after DMBA exposure, the mice were treated topically with twice-weekly doses of TPA (12-*O*-Tetradecanoylphorbol-13-acetate) in acetone (5 μg in 250 μL) for 15 weeks. The mice were euthanized 16 weeks after DMBA treatment. Papillomas were removed, and RNA was isolated by Tri Reagent extraction following the manufacturer’s protocol (Molecular Research Center; Cat.# TR118).

### Statistical Analysis

All statistical analysis was performed using GraphPad Prism version 8.3.1. The appropriate statistical test was used as indicated to determine p values of raw data. P-values < 0.05 were considered significant.

## Disclosure of Potential Conflicts of Interest

No potential conflicts of interest were disclosed.

## Authors’ Contributions

Conceptualization, K.E.L. and K.M.A.; Methodology, Y.I., A.V.K.; Investigation, K.E.L., R.B., E.S.D., C.W.C, E.L., Y.I., ad A.V.K.; Writing, K.E.L. and K.M.A.; Visualization, K.E.L., R.B., and K.M.A.; Supervision, Y.I., R.P.H., K.M.A.; Funding Acquisition, A.V.K., R.P.H., and K.M.A.

## Acknowledgements

We thank Dr. Naveen Kumar Tangudu for critical reading and editing of this manuscript. This work was supported by grants from the National Institutes of Health (F31CA250366 to K.E.L., F31CA236372 to E.S.D., and R00CA194309 and R37CA240625 to K.M.A.), the W. W. Smith Charitable Trust (to K.M.A.), and the Penn State Cancer Institute Postdoctoral Fellowship (to R.B.). Histone epiproteomics services were performed by the Northwestern Proteomics Core Facility, generously supported by NCI CCSG P30 CA060553 awarded to the Robert H Lurie Comprehensive Cancer Center, instrumentation award (S10OD025194) from NIH Office of Director, and the National Resource for Translational and Developmental Proteomics supported by P41 GM108569.

## Supplemental Figures

**Figure S1.**
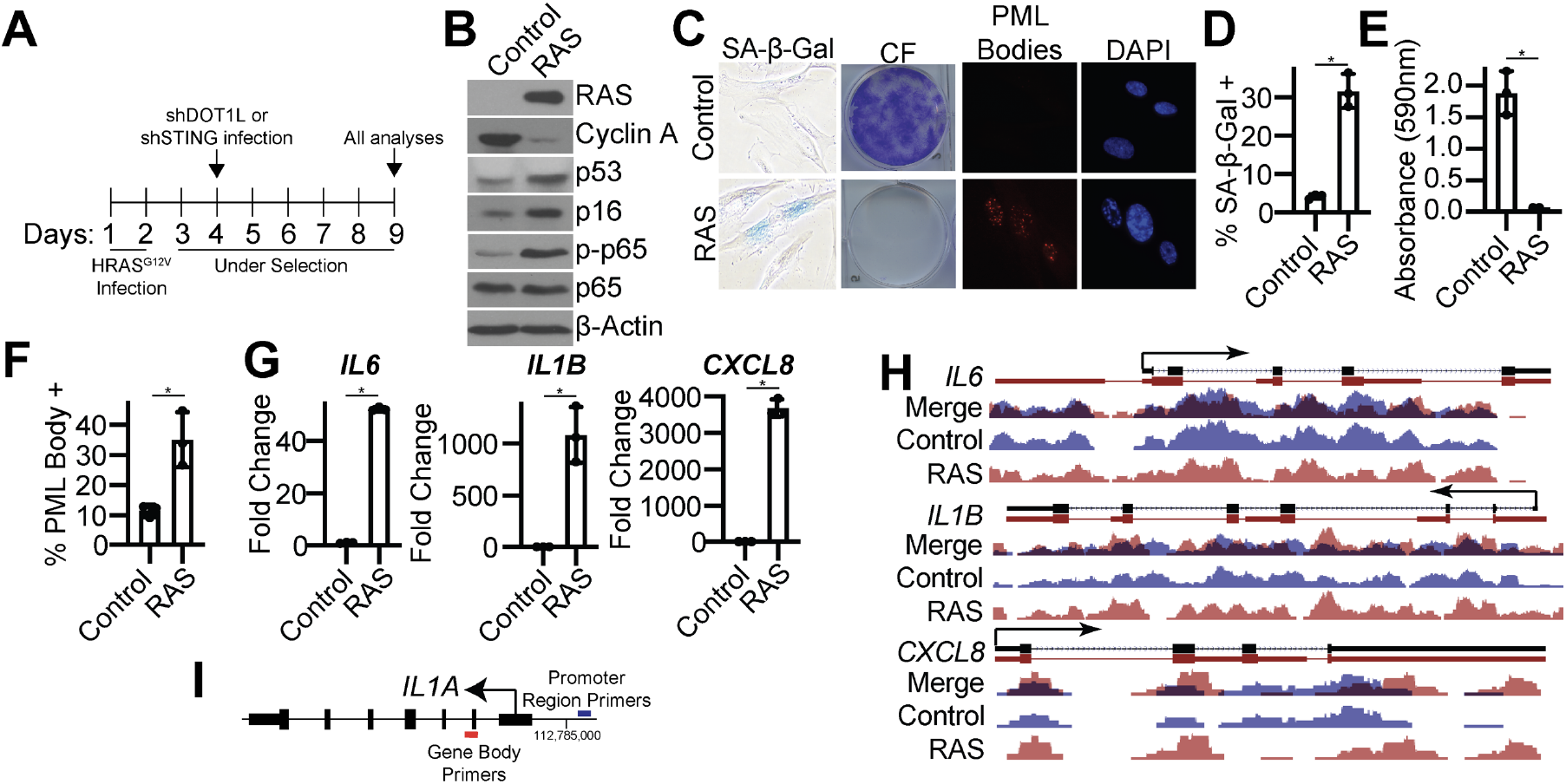
Oncogenic RAS induces cellular senescence; H3K79me3 is not enriched at other SASP genes. Related to Figure 1. **(A)** Timeline of experiments. **(B-H)** IMR90 cells were infected with retrovirus expressing HRas^G12V^ (RAS) or control. Cells were selected for 7 days with 1 μg/mL puromycin. **(B)** Immunoblot analysis of the indicated proteins. β-actin was used as a loading control. One of three experiments is shown. **(C)** SA-β-Gal activity, colony formation (CF), and PML body IF. One of three experiments is shown. **(D)** Quantification of SA-β-Gal activity in (C). Data represent mean ± SD. *p<0.0004. **(E)** Quantification of colony formation in (C). Data represent mean ± SD. *p<0.0008. **(F)** Quantification of PML body IF in (C). Data represent mean ± SD. *p<0.0118. **(G)** *IL6, IL1B*, and *CXCL8* mRNA expression. One of five experiments is shown. Data represent mean ± SD. *p<0.003. **(H)** H3K79me3 ChIP-Seq track at the *IL6, IL1B*, and *CXCL8* gene loci. Blue indicates binding of H3K79me3 in control cells, whereas red indicates binding of H3K79me3 binding in RAS cells. **(I)** Schematic of ChIP-qPCR primers for *IL1A* promoter region (H3K79me2 enrichment) and gene body (H3K79me3 enrichment).

**Figure S2.**
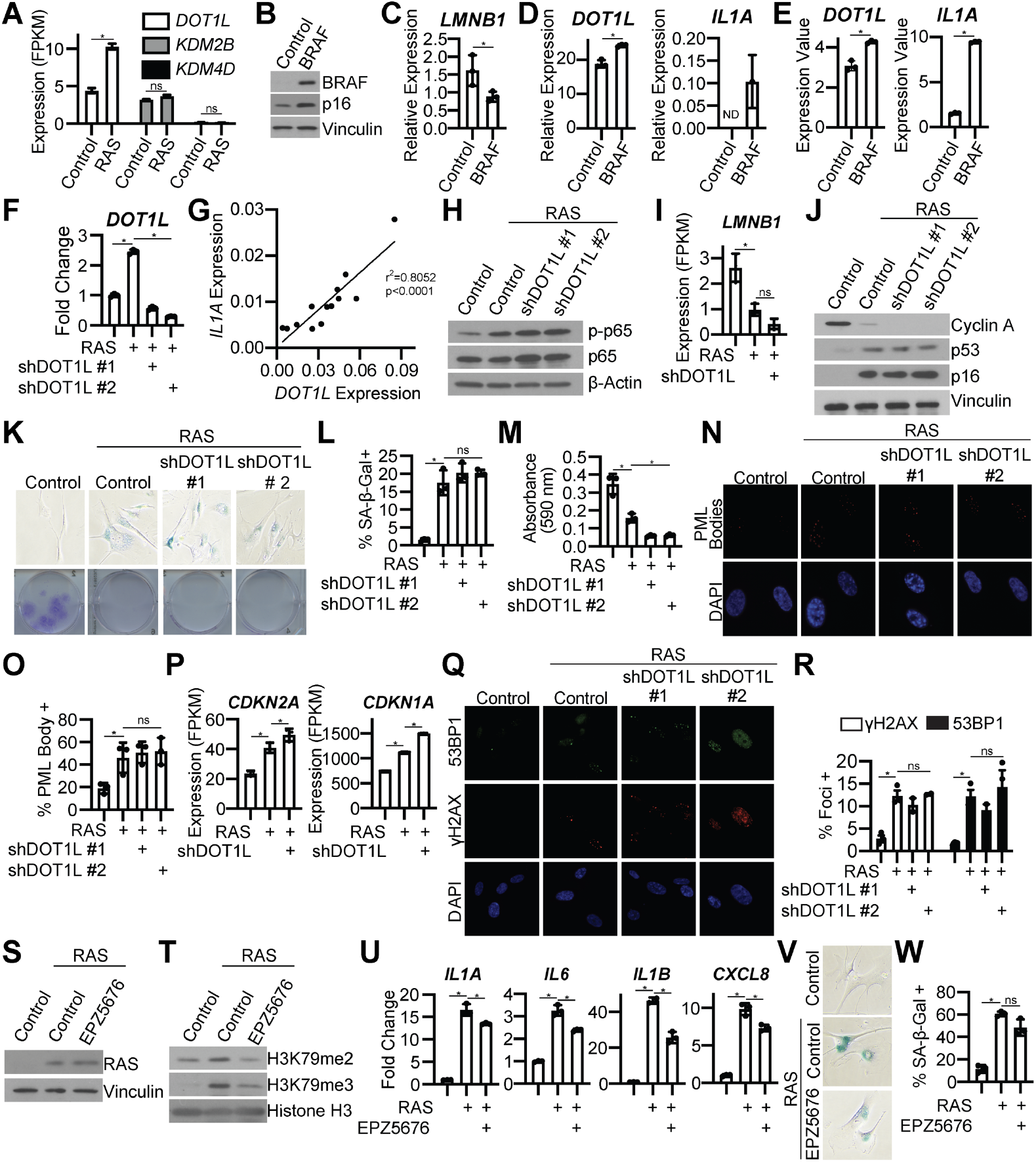
DOT1L knockdown maintains cells in a senescent-associated cell cycle arrest; DOT1L inhibitor decreases SASP gene expression. Related to Figure 2. **(A)** IMR90 cells were infected with retrovirus expressing HRAS^G12V^ (RAS) or control. *DOT1L, KDM2B*, and *KDM4D* mRNA expression from RNA-Seq. Data represent mean ± SD. *p<0.0001, ns=not significant. **(B-E)** Primary human melanocytes were infected with retrovirus expressing BRAF^V600E^ (BRAF) or control. Panel E are data from GSE46818. **(B)** BRAF and p16 immunoblot analysis. Vinculin was used as a loading control. One experiment is shown. **(C)** *LMNB1* mRNA expression was determined by RT-qPCR. One experiment is shown. Data represent mean ± SD. *p<0.05. **(D)** *DOT1L* and *IL1A* mRNA expression was determined by RT-qPCR. One experiment is shown. Data represent mean ± SD. ND=not detected. *p<0.005. **(E)** *DOT1L* and *IL1A e*xpression from microarray analysis (GSE46818). Data represent mean ± SD. *p<0.005. **(F and H-R)** IMR90 cells were infected with retrovirus expressing HRAS^G12V^ (RAS) or control with or without lentivirus expressing an shRNA to human DOT1L (shDOT1L). Details on timepoint can be found in Figure S1A. **(F)** *DOT1L* mRNA expression was determined by RT-qPCR. One of three experiments is shown. *p<0.0001. **(G)** RT-qPCR analysis was performed for *DOT1L* and *IL1A* expression on papillomas from mice treated with DMBA/TPA. **(H)** Immunoblot analysis of p-p65 and total p65. β-actin was used as a loading control. One of three experiments is shown. **(I)** *LMNB1* mRNA expression from RNA-Seq. Data represent mean ± SD. ns=not significant. *p<0.005. **(J)** Immunoblot analysis of cyclin A, p53, and p16. Vinculin was used as a loading control. One of three experiments is shown. **(K)** SA-β-Gal activity and colony formation. One of three experiments is shown. **(L)** Quantification of SA-β-Gal activity in (K). Data represent mean ± SD. *p<0.0014. **(M)** Quantification of colony formation in (K). Data represent mean ± SD. *p<0.05. **(N)** PML body IF. One of three experiments is shown. **(O)** Quantification of (N). Data represent mean ± SD. *p<0.02 vs control. **(P)** *CDKN2A* and *CDKN1A* mRNA expression from RNA-Seq. Data represent mean ± SD. *p<0.05. **(Q)** γH2AX and 53BP1 foci. One of three experiments is shown. **(R)** Quantification of (Q). Data represent mean ± SD. *p<0.05. **(S-W)** IMR90 cells were infected with retrovirus expressing HRAS^G12V^ (RAS) or control. Four days post-retroviral infection cells were treated with 1 μM DOT1L inhibitor EPZ5676. **(S)** RAS immunoblot analysis. Vinculin was used as loading control. One of three experiments is shown. **(T)** H3K79me2 and H3K79me3 immunoblot analysis was performed on chromatin fractions. Total histone H3 was used as loading control. One of three experiments is shown. **(U)** *IL1A, IL6, IL1B*, and *CXCL8* mRNA expression was determined by RT-qPCR. One of three experiments is shown. Data represent mean ± SD. *p<0.01. **(V)** SA-β-Gal activity. One of three experiments is shown. **(W)** Quantification of SA-β-Gal activity in (V). Data represent mean ± SD. *p<0.0005.

**Figure S3.**
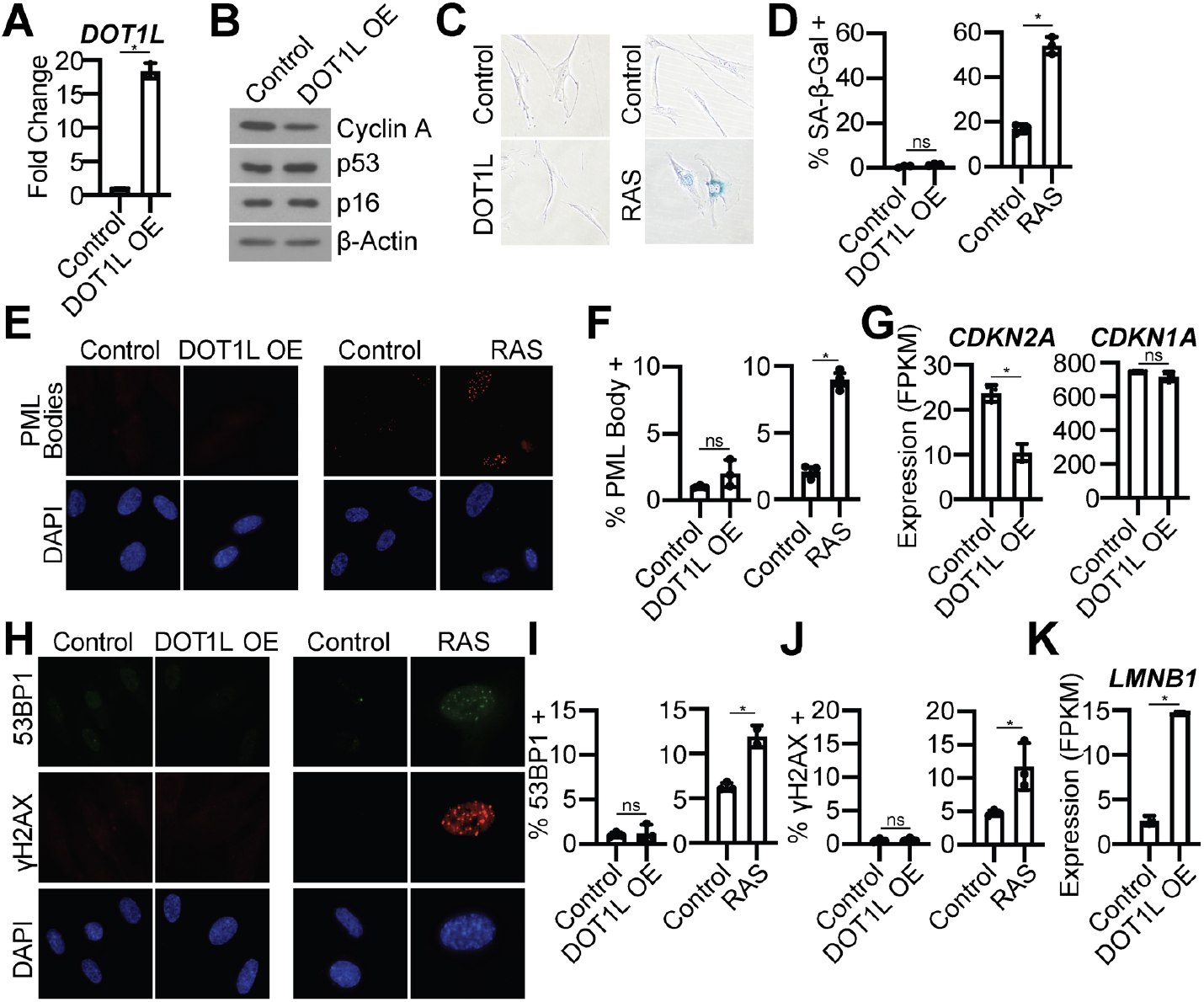
DOT1L overexpression does not induce a senescent-associated cell cycle arrest or DNA damage accumulation. Related to Figure 3. **(A-K)** IMR90 cells were infected with retrovirus expressing human DOT1L or control. Cells were selected for 4 days with 1 μg/mL blasticidin. As a positive control, IMR90 cells were infected with retrovirus expressing HRAS^G12V^ (RAS) or control. Cells were selected for 7 days with 1 μg/mL puromycin. **(A)** *DOT1L* mRNA expression. One of three experiments is shown. Data represent mean ± SD. *p<0.001. **(B)** Immunoblot analysis of the indicated proteins. β-actin was used as a loading control. One of three experiments is shown. **(C)** SA-β-Gal activity. One of three experiment is shown. **(D)** Quantification of (C). Data represent mean ± SD. ns=not significant, *p<0.001. **(E)** PML body IF. One of three experiments is shown. **(F)** Quantification of (E). Data represent mean ± SD. ns=not significant, *p<0.01. **(G)** *CDKN2A* and *CDKN1A* mRNA expression from RNA-Seq. Data represent mean ± SD. *p<0.005. **(H)** γH2AX and 53BP1 IF. One of three experiments is shown. **(I)** Quantification of 53BP1 foci in (H). Data represent mean ± SD. ns=not significant, *p<0.005. **(J)** Quantification of γH2AX foci in (H). Data represent mean ± SD. ns=not significant, *p<0.05. **(K)** *LMNB1* expression from RNA-Seq. Data represent mean ± SD. *p<0.0001.

**Figure S4.**
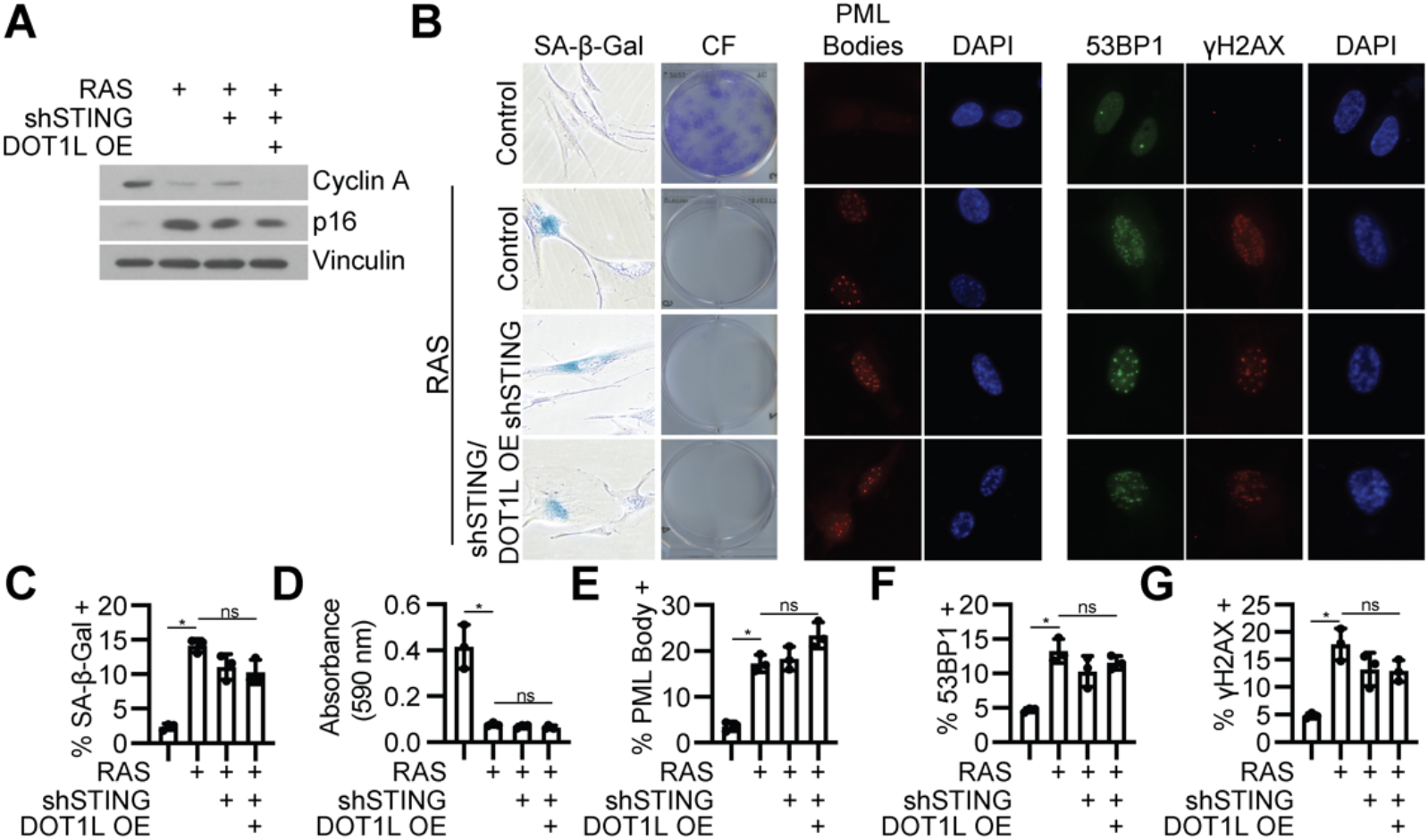
STING knockdown and DOT1L overexpressing maintains the senescence-associated cell cycle arrest. Related to Figure 4. **(A-G)** IMR90 cells were infected with retrovirus expressing HRas^G12V^ (RAS) or control with or without lentivirus expressing an shRNA to human STING (shSTING) with or without overexpression of DOT1L. (A) Immunoblot analysis for the indicated proteins. Vinculin was used as a loading control. One of three experiments is shown. **(B)** SA-β-Gal activity, colony formation (CF), PML body IF, and γH2AX and 53BP1 IF. One of three experiment is shown. **(C)** Quantification of SA-β-Gal in (B). Data represent mean ± SD. *p<0.01. ns=not significant. **(D)** Quantification of colony formation in (B). Data represent mean ± SD. *p<0.01. ns=not significant. **(E)** Quantification of PML body foci in (B). Data represent mean ± SD. *p<0.02 vs control. ns=not significant. **(F)** Quantification of 53BP1 foci in (B). Data represent mean ± SD. *p<0.01 vs control. ns=not significant. **(G)** Quantification of γH2AX foci in (B). Data represent mean ± SD. *p<0.01 vs control. ns=not significant.

## Supplemental Tables

**Table S1:** Epiproteomics analysis in RAS vs. Control cells

**Table S2:** RAS vs. control genes with peaks (FC>1.5) in ChIP-Seq and upregulated (FC>1.5; FDR<0.25) in RNA-Seq

**Table S3:** Integrated density of cytokines from antibody array.

**Table S4:** Primers used for the studies.

**Table S5:** Annotated ChIP-Seq peaks and analysis

**Table S6:** RNA-Seq differential expression analysis

## Notes

### Competing Interest Statement

The authors have declared no competing interest.

